# syntenet: an R/Bioconductor package for the inference and analysis of synteny networks

**DOI:** 10.1101/2022.08.16.504079

**Authors:** Fabricio Almeida-Silva, Tao Zhao, Kristian K. Ullrich, M. Eric Schranz, Yves Van de Peer

## Abstract

Interpreting and visualizing synteny relationships across several genomes is a challenging task. We previously proposed a network-based approach for better visualization and interpretation of large-scale microsynteny analyses. Here, we present *syntenet*, an R package to infer and analyze synteny networks from whole-genome protein sequence data. The package offers a simple and complete framework, including data preprocessing, synteny detection and network inference, network clustering and phylogenomic profiling, and microsynteny-based phylogeny inference. Graphical functions are also available to create publication-ready plots. Synteny networks inferred with *syntenet* can highlight taxon-specific gene clusters that likely contributed to the evolution of important traits, and microsynteny-based phylogenies can help resolve phylogenetic relationships under debate.

**Availability and Implementation:** The package is available on Bioconductor (https://bioconductor.org/packages/syntenet), and the source code is available on a GitHub repository (https://github.com/almeidasilvaf/syntenet).

## 1 Introduction

Gene and genome duplications provide organisms with the raw genetic material for biological innovations (Ohno, 1970; Panchy *et al*., 2016; Van De Peer *et al*., 2017). Thus, exploring the evolution of duplicated genes and genomes can help explain how new traits arise and diversify across taxa. Identifying collinear or syntenic regions (here used as synonyms, i.e., different genomic segments showing conserved gene content and order) within genomes has become standard practice to detect signatures of whole-genome duplications (WGD) and the genomic rearrangements that typically follow WGD events (Liu *et al*., 2022; Ma *et al*., 2021; Vanneste *et al*., 2013; Wan *et al*., 2021). Synteny analyses can also be performed to compare different genomes to provide insights on population structure, species divergence, and the evolution of gene families and traits (Jayakodi *et al*., 2020; Li *et al*., 2022; Tang *et al*., 2022; Zhang *et al*., 2021; Zhou *et al*., 2017). However, when comparing synteny relationships among several genomes, interpretation and visualization is notoriously complex.

We previously proposed a network-based approach to analyze synteny in large datasets that consists in treating anchor pairs (duplicates retained from a large-scale duplication event) from synteny comparisons as connected nodes of an undirected unweighted graph (Zhao and Schranz, 2017). We have used such synteny networks to study the evolution of MADS-box transcription factors in plants (Zhao *et al*., 2017), explore the conservation patterns of synteny clusters in mammalian and angiosperm genomes (Zhao and Schranz, 2019), and to provide insights into controversial phylogenetic relationships in angiosperms through a microsynteny-based phylogeny (Zhao *et al*., 2021). However, despite gaining wide interest, and its wide applicability, our method has not been incorporated in a distributable format. Here, we present *syntenet*, an R/Bioconductor package to infer and analyze synteny networks from whole-genome protein sequences. *syntenet* integrates our whole previously developed method in an easy and simple framework, from data preprocessing to network analyses, and visualization.

## 2 Implementation

For seamless integration with other Bioconductor packages, the input objects for *syntenet* are base R or core Bioconductor classes (Fig. 1A). The complete pipeline requires the external software tools DIAMOND (Buchfink *et al*., 2021) and IQTREE2 (Minh *et al*., 2020). Users must input: i. protein sequences for each species, stored in a list of AAStringSet objects and; ii. genomic coordinates for each gene, stored in a GRangesList object.

**Fig. 1.**
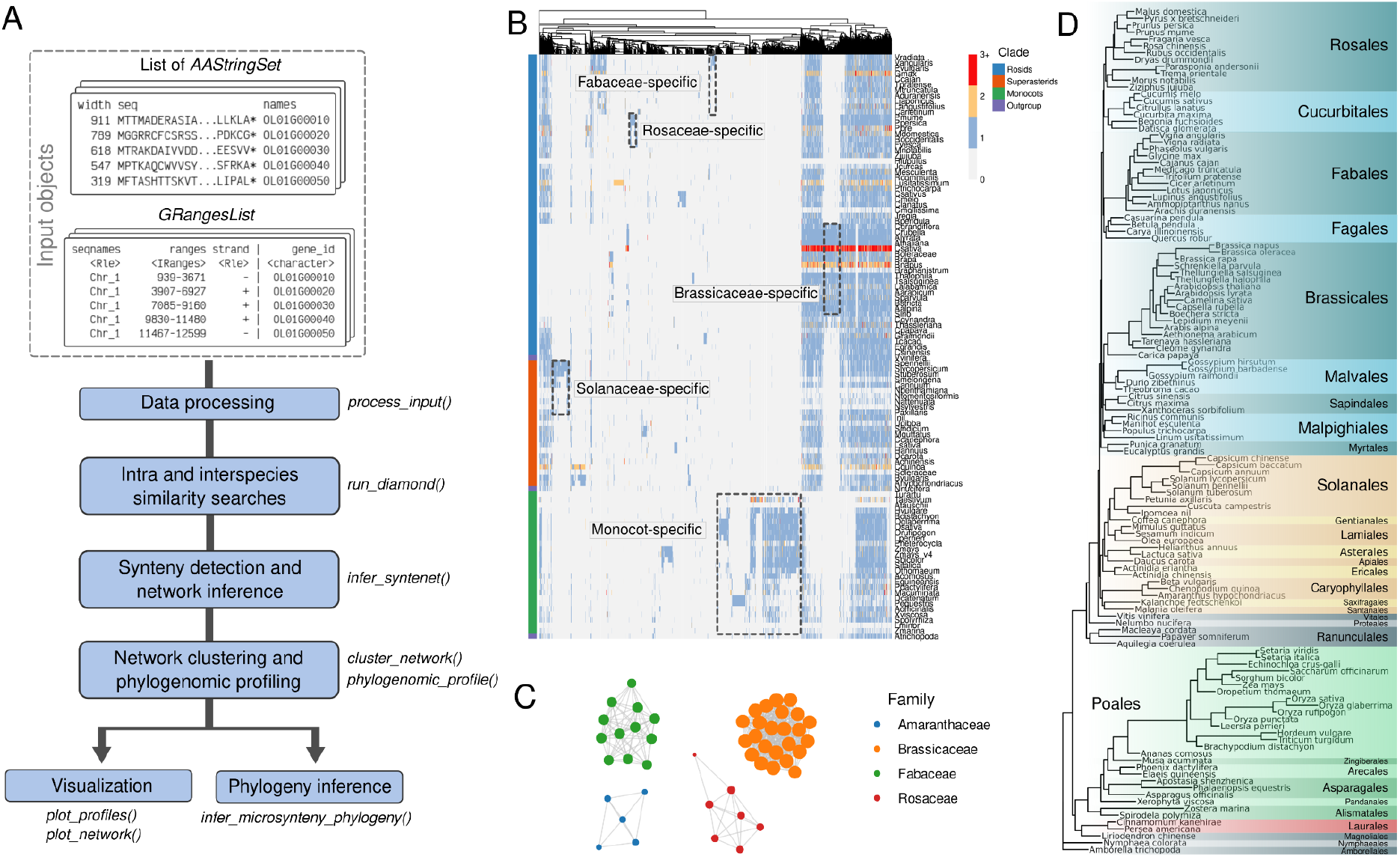
Workflow and possible applications of syntenet. A. Schematic model of the whole pipeline for synteny network inference and analysis. Blue boxes represent the different steps of the pipeline, with the corresponding function names in italicized fonts on the right and below. B. Example heatmap of phylogenomic profiles that can be created with the function *phylogenomic_profile()*. Dashed boxes highlight family-specific synteny clusters. Interestingly, species with recent whole-genome duplications (e.g., *Malus domestica* and *Glycine max*) have a higher number of genes in most synteny clusters. Data were obtained from (Zhao and Schranz, 2019), and code to create the figure is available on Supplementary Data S1. C. Network representation of some family-specific synteny clusters from Fig. 1B produced with the function *plot_network*. D. Microsynteny-based phylogeny of angiosperms. Data were obtained from (Zhao *et al*., 2021), and code to reproduce the figure is available on Supplementary Data S2.

### 2.1 Data preprocessing and similarity search

The input data is automatically preprocessed to clean up gene names, add unique species identifiers to gene and chromosome names, and to make sure that only translated sequences from primary transcripts are included. The processed data is used as input to sequence similarity search programs, such as BLASTp (Altschul *et al*., 1997) or DIAMOND (Buchfink *et al*., 2021). Although users can run DIAMOND or BLAST on the command line, *syntenet* has a wrapper function named *run_diamond* that runs DIAMOND from the R session and reads the tabular output as a data frame (Fig. 1A).

### 2.2 Synteny detection

The function *infer_syntenet* integrates gene coordinates with the DIAMOND output to detect anchor pairs (Fig. 1A). Synteny detection is performed using a native version of the popular MCScanX algorithm (Wang *et al*., 2012), which has been ported to R with the Rcpp framework to integrate C++ code in R packages (Eddelbuettel and François, 2011). Hence, users have access to the same accuracy and speed of MCScanX without having to install it. It is noteworthy that synteny detection in R can also be performed with the Bioconductor package DECIPHER (Wright, 2016), which has its own synteny detection algorithm, but whose accuracy has not been benchmarked against existing tools.

### 2.3 Synteny network clustering and phylogenomic profiling

The synteny network inferred with *infer_syntenet* is represented as an edge list. Network clustering is performed with the Infomap algorithm, which has been demonstrated as the best clustering technique for synteny networks (Zhao *et al*., 2021). Synteny clusters are used for phylogenomic profiling, which consists in obtaining a matrix m_ij_ representing the number of genes from cluster *j* that can be found in species *i* (Fig. 1B). Phylogenomic profiles are clustered using Ward’s clustering on a matrix of Jaccard distances. This analysis can reveal synteny clusters that are deeply conserved across taxa, and taxon-specific clusters (*e*.*g*., family-specific synteny clusters in Fig. 1B). Clusters can be visualized either as a heatmap (Fig. 1B) or as a network plot (Fig. 1C).

### 2.4 Microsynteny-based phylogeny reconstruction

For phylogeny inference, the matrix of phylogenomic profiles is binarized, transposed, and exported as a PHYLIP-formatted file. The function *infer_microsynteny_phylogeny* passes this PHYLIP file to IQTREE2, which infers a phylogeny from binary data using the MK+FO+R model with 1000 bootstrap replicates and 1000 replicates for the SH-like approximate likelihood ratio test. Users can also choose a different substitution model, provided that it is suitable for binary data.

## 3 Application to real data sets

We demonstrated the effectiveness of *syntenet* by reproducing results from previous works on synteny networks. We recreated phylogenomic profiles for angiosperm genomes from (Zhao and Schranz, 2019), and we reconstructed the microsynteny-based phylogeny of angiosperms from (Zhao *et al*., 2021) (Supplementary Data S1 and S2). We have also inferred synteny networks from 16 algae genomes (phylum Chlorophyta) available on Pico-PLAZA 3.0 (Van Bel *et al*., 2018). Synteny detection and network inference for 17 genomes (16 Chlorophyta and *Physcomitrium patens* as outgroup) took ∼55 seconds on an Ubuntu 20.04 laptop with an Intel i5-1135G7 processor (2.40 GHz; 8 GB RAM) (Supplementary Data S3).

## 4 Known limitation

Synteny detection in highly fragmented genomes is challenging, so the MCScanX algorithm might fail to detect some syntenic blocks in such genomes (Liu *et al*., 2018). Thus, when selecting species to use in *syntenet*, we recommend using genomes with at least 85% complete BUSCOs to avoid bias. However, with the fast advancement in sequencing technologies and resequencing of low-quality reference genomes, genome completeness will likely cease to be a concern in the near future.

## 5 Conclusions

*syntenet* is an R package to infer synteny networks from whole-genome protein sequence data. Synteny networks can be explored to detect deeply conserved and taxa-specific clusters, to explore genomic rearrangements, to better visualize syntenic relationships, and to infer microsynteny-based phylogenies. The user-friendly framework in *syntenet* will likely make synteny networks widely used in comparative genomics.

## Acknowledgements

YVdP acknowledges funding from the European Research Council (ERC) under the European Union’s Horizon 2020 research and innovation program (No. 833522). YVdP and FA-S acknowledge funding from Ghent University (Methusalem funding, BOF.MET.2021.0005.01). KKU acknowledges institutional funding through the Max Planck Society. TZ acknowledges the Chinese Universities Scientific Fund 2452021133.

## Conflicts of interest

none declared.

## Notes

### Competing Interest Statement

The authors have declared no competing interest.

### Summary of Updates

This version of the manuscript has been revised to update the authors list

