## Supplementary Text for "syntenet: an R/Bioconductor package for the inference and analysis of synteny networks"

**19 August 2022**

### Introduction

Here, we will use `syntenet` to reproduce the findings from two of our previous papers on synteny networks:

1. Zhao, T., & Schranz, M. E. (2019). Network-based microsynteny analysis identifies major differences and genomic outliers in mammalian and angiosperm genomes. **Proceedings of the National Academy of Sciences**, 116(6), 2165-2174. DOI: 10.1073/pnas.1801757116
2. Zhao, T., Zwaenepoel, A., Xue, J. Y., Kao, S. M., Li, Z., Schranz, M. E., & Van de Peer, Y. (2021). Whole-genome microsynteny-based phylogeny of angiosperms. **Nature Communications**, 12(1), 1-14. DOI: 10.1038/s41467-021-23665-0

Besides, we will run the whole pipeline on a new data set of algae genomes from the Chlorophyta clade. The data were obtained from Pico-PLAZA 3.0 (Van Bel et al. 2018).

```
library(syntenet)
library(tidyverse)
set.seed(123)
```

### Section 1: Recreating phylogenomic profiles from Zhao and Schranz (2019)

In this section, we will identify phylogenomic profiles from synteny clusters of BUSCO genes in angiosperms. Then, we will represent the phylogenomic profiles as in Figure 1B of the manuscript. The data for this section were obtained from the original publication (Zhao and Schranz 2019), in its Dataverse repository (DOI: 10.7910/DVN/BDMA7A).

```
#----Download file-------------------------------------------------------------
net_file <- file.path(tempdir(), "busco_network.txt.gz")
if(!file.exists(net_file)) {
    download.file("https://dataverse.harvard.edu/api/access/datafile/:persistentId?persistentId=doi:10.7910/I
}

#----Read file-----------------------------------------------------------------
busco_network <- readr::read_delim(net_file, show_col_types = FALSE,
                                   col_names = FALSE, delim = " ")

## Create a data frame of taxonomic information for each species
### Family and abbreviations
species_fam <- data.frame(
    Species = c(
        "Vigna radiata", "Vigna angularis", "Phaseolus vulgaris",
        "Glycine max", "Cajanus cajan", "Trifolium pratense",
        "Medicago truncatula", "Arachis duranensis", "Lotus japonicus",
        "Lupinus angustifolius", "Cicer arietinum",
        "Prunus mume", "Prunus persica", "Pyrus x bretschneideri",
        "Malus domestica", "Rubus occidentalis", "Fragaria vesca",
        "Morus notabilis", "Ziziphus jujuba", "Humulus lupulus",
```

```
        "Jatropha curcas", "Manihot esculenta", "Ricinus communis",
        "Linum usitatissimum", "Populus trichocarpa", "Cucumus sativus",
        "Cucumis melo", "Citrullus lanatus",
        "Castanea mollissima", "Juglans regia", "Betula pendula",
        "Capsella grandiflora", "Capsella rubella", "Arabidopsis lyrata",
        "Arabidopsis thaliana", "Camelina sativa", "Brassica oleraceae",
        "Brassica rapa", "Brassica napus", "Raphanus raphanistrum",
        "Thellungiella halophila", "Thellungiella salsuginea",
        "Leavenworthia alabamica", "Aethionema arabicum",
        "Schrenkiella parvula", "Boechera stricta", "Arabis alpina",
        "Sisymbrium irio", "Cleome gynandra", "Tarenaya hassleriana",
        "Carica papaya", "Gossypium raimondii", "Theobroma cacao",
        "Eucalyptus grandis", "Citrus sinensis", "Vitis vinifera",
        "Solanum pennellii", "Solanum lycopersicum", "Solanum tuberosum",
        "Solanum melongena", "Capsicum annuum", "Nicotiana benthamiana",
        "Nicotiana tomentosiformis", "Nicotiana attenuata",
        "Nicotiana sylvestris", "Petunia axillaris",
        "Ipomoea nil", "Utricularis gibba", "Sesamum indicum",
        "Mimulus guttatus", "Coffea canephora", "Lactuca sativa",
        "Helianthus annuus", "Daucus carota",
        "Actinidia chinensis", "Chenopodium quinoa", "Spinacia oleraceae",
        "Beta vulgaris", "Amaranthus hypochondriacus",
        "Nelumbo nucifera", "Triticum urartu", "Triticum aestivum",
        "Aegilops tauschii", "Hordeum vulgare", "Brachypodium distachyon",
        "Oryza glaberrima", "Oryza sativa", "Oryza rufipogon",
        "Leersia perrieri", "Phylostachys heterocycla", "Zea mays",
        "Zea mays V4", "Sorghum bicolor", "Setaria italica",
        "Oropetium thomaeum", "Ananas comosus", "Elaeis guineensis",
        "Phoenix dactylifera", "Musa acuminata", "Dendrobium catenatum",
        "Phalaenopsis equestris", "Asparagus officinalis", "Xerophyta viscosa",
        "Spirodela polyrhiza", "Lemna minor", "Zostera marina",
        "Amborella trichopoda"
    ),
    Abbrev = c(
        "vra", "van", "pvu", "gma", "cca", "tpr", "mtr", "adu", "lja", "Lang",
        "car", "pmu", "ppe", "pbr", "mdo", "roc", "fve", "Mnot", "Zjuj",
        "hlu", "jcu", "mes", "rco", "lus", "ptr", "csa", "cme", "cla",
        "cmo", "jre", "Bpen", "cgr", "cru", "Alyr", "ath", "Csat", "bol",
        "bra", "bnp", "rra", "thh", "tsa", "lal", "aar", "spa", "Bostr", "Alp",
        "sir", "cgy", "tha", "cpa", "gra", "tca", "egr", "csi", "vvi",
        "spe", "sly", "stu", "sme", "can", "nbe", "Ntom", "Natt",
        "Nsyl", "pax", "Inil", "ugi", "sin", "mgu", "coc", "Lsat", "hel",
        "dca", "ach", "Cqui", "sol", "bvu", "Ahyp", "nnu", "tur", "tae", "ata",
        "HORVU", "bdi", "ogl", "osa", "oru", "lpe", "phe", "zma", "Zmay",
        "sbi", "sit", "oth", "aco", "egu", "Pdac", "mac", "Dcat", "peq",
        "Aoff", "Xvis", "spo", "lmi", "zom", "atr"
    ),
    Family = c(
        rep("Fabaceae", 11), rep("Rosaceae", 6),
        "Moraceae", "Rhamnaceae", "Cannabaceae",
        rep("Euphorbiaceae", 3), "Linaceae", "Salicaceae",
```

```
        rep("Cucurbitaceae", 3), "Fagaceae", "Juglandaceae", "Betulaceae",
        rep("Brassicaceae", 17), rep("Cleomaceae", 2), "Caricaceae",
        rep("Malvaceae", 2), "Myrtaceae", "Rutaceae", "Vitaceae",
        rep("Solanaceae", 10), "Convolvulaceae", "Lentibulariaceae",
        "Pedaliaceae", "Phrymaceae", "Rubiaceae", rep("Asteraceae", 2),
        "Apiaceae", "Actinidiaceae", rep("Amaranthaceae", 4), "Nelumbonaceae",
        rep("Poaceae", 15), "Bromeliaceae", rep("Arecaceae", 2), "Musaceae",
        rep("Orchidaceae", 2), "Asparagaceae", "Velloziaceae",
        rep("Araceae", 2), "Zosteraceae", "Amborellaceae"
    )
)

### Superorders
species_sorder <- data.frame(
    Family = unique(species_fam$Family)
)

### Superorders
species_sorder$Clade <- c(
    rep("Rosids", 18), "Outgroup", rep("Superasterids", 10), "Outgroup",
    rep("Monocots", 9), "Outgroup"
)

### All in one
species_tax <- merge(species_fam, species_sorder, by = "Family", sort = FALSE)
species_tax <- species_tax[, c("Species", "Abbrev", "Family", "Clade")]
species_tax # inspect the data
##                      Species Abbrev           Family         Clade
## 1              Vigna radiata    vra         Fabaceae        Rosids
## 2            Vigna angularis    van         Fabaceae        Rosids
## 3         Phaseolus vulgaris    pvu         Fabaceae        Rosids
## 4                Glycine max    gma         Fabaceae        Rosids
## 5              Cajanus cajan    cca         Fabaceae        Rosids
## 6         Trifolium pratense    tpr         Fabaceae        Rosids
## 7        Medicago truncatula    mtr         Fabaceae        Rosids
## 8         Arachis duranensis    adu         Fabaceae        Rosids
## 9            Lotus japonicus    lja         Fabaceae        Rosids
## 10      Lupinus angustifolius   Lang         Fabaceae        Rosids
## 11            Cicer arietinum    car         Fabaceae        Rosids
## 12                Prunus mume    pmu         Rosaceae        Rosids
## 13             Prunus persica    ppe         Rosaceae        Rosids
## 14      Pyrus x bretschneideri    pbr         Rosaceae        Rosids
## 15            Malus domestica    mdo         Rosaceae        Rosids
## 16         Rubus occidentalis    roc         Rosaceae        Rosids
## 17             Fragaria vesca    fve         Rosaceae        Rosids
## 18            Morus notabilis   Mnot         Moraceae        Rosids
## 19            Ziziphus jujuba   Zjuj       Rhamnaceae        Rosids
## 20            Humulus lupulus    hlu      Cannabaceae        Rosids
## 21            Jatropha curcas    jcu    Euphorbiaceae        Rosids
## 22          Manihot esculenta    mes    Euphorbiaceae        Rosids
## 23            Ricinus communis    rco    Euphorbiaceae        Rosids
```

```
## 24        Linum usitatissimum    lus        Linaceae        Rosids
## 25        Populus trichocarpa    ptr      Salicaceae        Rosids
## 26            Cucumus sativus    csa   Cucurbitaceae        Rosids
## 27               Cucumis melo    cme   Cucurbitaceae        Rosids
## 28          Citrullus lanatus    cla   Cucurbitaceae        Rosids
## 29        Castanea mollissima    cmo        Fagaceae        Rosids
## 30              Juglans regia    jre    Juglandaceae        Rosids
## 31             Betula pendula   Bpen      Betulaceae        Rosids
## 32       Capsella grandiflora    cgr    Brassicaceae        Rosids
## 33           Capsella rubella    cru    Brassicaceae        Rosids
## 34         Arabidopsis lyrata   Alyr    Brassicaceae        Rosids
## 35       Arabidopsis thaliana    ath    Brassicaceae        Rosids
## 36            Camelina sativa   Csat    Brassicaceae        Rosids
## 37         Brassica oleraceae    bol    Brassicaceae        Rosids
## 38              Brassica rapa    bra    Brassicaceae        Rosids
## 39             Brassica napus    bnp    Brassicaceae        Rosids
## 40      Raphanus raphanistrum    rra    Brassicaceae        Rosids
## 41    Thellungiella halophila    thh    Brassicaceae        Rosids
## 42   Thellungiella salsuginea    tsa    Brassicaceae        Rosids
## 43    Leavenworthia alabamica    lal    Brassicaceae        Rosids
## 44        Aethionema arabicum    aar    Brassicaceae        Rosids
## 45       Schrenkiella parvula    spa    Brassicaceae        Rosids
## 46           Boechera stricta  Bostr    Brassicaceae        Rosids
## 47              Arabis alpina    Alp    Brassicaceae        Rosids
## 48            Sisymbrium irio    sir    Brassicaceae        Rosids
## 49            Cleome gynandra    cgy      Cleomaceae        Rosids
## 50       Tarenaya hassleriana    tha      Cleomaceae        Rosids
## 51              Carica papaya    cpa      Caricaceae        Rosids
## 52        Gossypium raimondii    gra       Malvaceae        Rosids
## 53            Theobroma cacao    tca       Malvaceae        Rosids
## 54         Eucalyptus grandis    egr       Myrtaceae        Rosids
## 55            Citrus sinensis    csi        Rutaceae        Rosids
## 56             Vitis vinifera    vvi        Vitaceae      Outgroup
## 57          Solanum pennellii    spe      Solanaceae Superasterids
## 58       Solanum lycopersicum    sly      Solanaceae Superasterids
## 59          Solanum tuberosum    stu      Solanaceae Superasterids
## 60          Solanum melongena    sme      Solanaceae Superasterids
## 61            Capsicum annuum    can      Solanaceae Superasterids
## 62      Nicotiana benthamiana    nbe      Solanaceae Superasterids
## 63  Nicotiana tomentosiformis   Ntom      Solanaceae Superasterids
## 64        Nicotiana attenuata   Natt      Solanaceae Superasterids
## 65       Nicotiana sylvestris   Nsyl      Solanaceae Superasterids
## 66          Petunia axillaris    pax      Solanaceae Superasterids
## 67               Ipomoea nil   Inil  Convolvulaceae Superasterids
## 68          Utricularis gibba    ugi Lentibulariaceae Superasterids
## 69            Sesamum indicum    sin     Pedaliaceae Superasterids
## 70           Mimulus guttatus    mgu     Phrymaceae Superasterids
## 71           Coffea canephora    coc       Rubiaceae Superasterids
## 72             Lactuca sativa   Lsat      Asteraceae Superasterids
## 73          Helianthus annuus    hel      Asteraceae Superasterids
## 74               Daucus carota    dca       Apiaceae Superasterids
```

```
## 75        Actinidia chinensis    ach    Actinidiaceae Superasterids
## 76         Chenopodium quinoa   Cqui    Amaranthaceae Superasterids
## 77         Spinacia oleraceae    sol    Amaranthaceae Superasterids
## 78              Beta vulgaris    bvu    Amaranthaceae Superasterids
## 79  Amaranthus hypochondriacus   Ahyp    Amaranthaceae Superasterids
## 80            Nelumbo nucifera    nnu    Nelumbonaceae      Outgroup
## 81             Triticum urartu    tur          Poaceae      Monocots
## 82           Triticum aestivum    tae          Poaceae      Monocots
## 83           Aegilops tauschii    ata          Poaceae      Monocots
## 84             Hordeum vulgare  HORVU          Poaceae      Monocots
## 85      Brachypodium distachyon    bdi          Poaceae      Monocots
## 86           Oryza glaberrima    ogl          Poaceae      Monocots
## 87               Oryza sativa    osa          Poaceae      Monocots
## 88             Oryza rufipogon    oru          Poaceae      Monocots
## 89            Leersia perrieri    lpe          Poaceae      Monocots
## 90     Phylostachys heterocycla    phe          Poaceae      Monocots
## 91                    Zea mays    zma          Poaceae      Monocots
## 92                 Zea mays V4   Zmay          Poaceae      Monocots
## 93             Sorghum bicolor    sbi          Poaceae      Monocots
## 94             Setaria italica    sit          Poaceae      Monocots
## 95          Oropetium thomaeum    oth          Poaceae      Monocots
## 96              Ananas comosus    aco     Bromeliaceae      Monocots
## 97           Elaeis guineensis    egu        Arecaceae      Monocots
## 98         Phoenix dactylifera   Pdac        Arecaceae      Monocots
## 99              Musa acuminata    mac         Musaceae      Monocots
## 100        Dendrobium catenatum   Dcat      Orchidaceae      Monocots
## 101      Phalaenopsis equestris    peq      Orchidaceae      Monocots
## 102       Asparagus officinalis   Aoff     Asparagaceae      Monocots
## 103          Xerophyta viscosa   Xvis     Velloziaceae      Monocots
## 104        Spirodela polyrhiza    spo          Araceae      Monocots
## 105                Lemna minor    lmi          Araceae      Monocots
## 106             Zostera marina    zom      Zosteraceae      Monocots
## 107         Amborella trichopoda    atr    Amborellaceae      Outgroup

#----Add underscore (i.e., "_") in front of species abbreviations in genes------
new_abbrev <- paste0(species_fam$Abbrev, "_")
names(new_abbrev) <- paste0("^", species_fam$Abbrev)

busco_network <- busco_network %>%
    dplyr::mutate(X1 = str_replace_all(X1, new_abbrev)) %>%
    dplyr::mutate(X1 = str_replace_all(X1, "__", "_")) %>%
    dplyr::mutate(X2 = str_replace_all(X2, new_abbrev)) %>%
    dplyr::mutate(X2 = str_replace_all(X2, "__", "_"))

#----Cluster network and get phylogenomic profiles-------------------------------
clusters <- cluster_network(busco_network)
profiles <- phylogenomic_profile(clusters)

## Rename species in profiles matrix with:
## First letter of genus, whole specific epithet (e.g., Athaliana, Gmax, etc.)
new_tax <- species_tax %>%
```

```
    mutate(Genus = word(Species)) %>%
    mutate(Genus = str_sub(Genus, 1, 1)) %>%
    mutate(se = word(Species, 2)) %>%
    mutate(names = str_c(Genus, se))
new_tax$names[new_tax$names == "Px"] <- "Pbre"
new_tax$names[92] <- "Zmays_v4"

## Get species order for the heatmap
heatmap_order <- new_tax$names
new_tax <- new_tax %>%
    arrange(Abbrev)
colnames(profiles$profile_matrix) <- new_tax$names

#----Plot profiles--------------------------------------------------------------
plot_profiles(
    profiles,
    species_annotation = new_tax[, c("names", "Clade")],
    cluster_species = heatmap_order,
    cluster_columns = TRUE
)
```

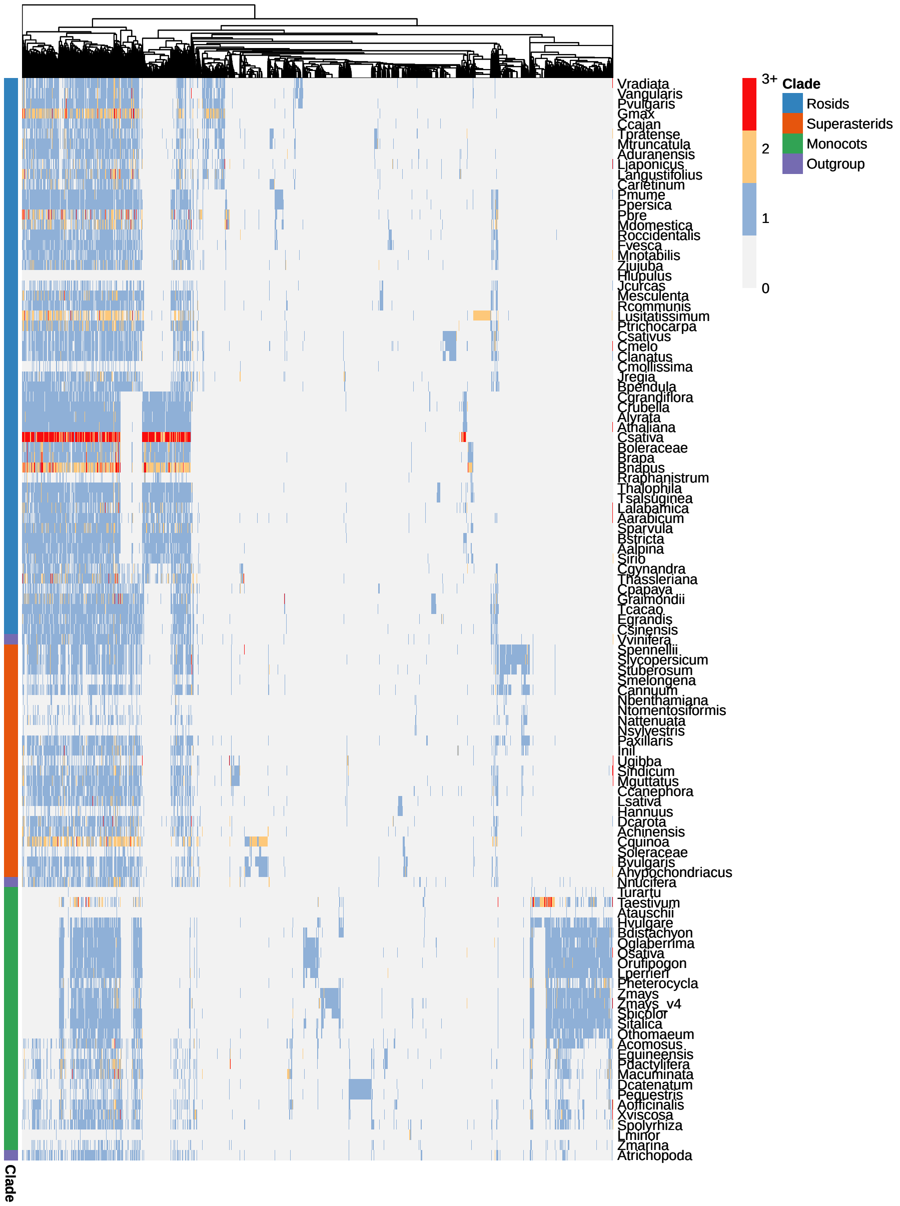

As in Figure 1B, we can see deeply conserved synteny clusters and clade-specific clusters.

### Section 2: Rebuilding the angiosperm phylogeny from Zhao et al. (2021)

Here, we will infer a microsynteny-based phylogeny of angiosperms, as we did in Zhao et al. (2021). Data were obtained from the Dataverse repository associated with the original publication (DOI: 10.7910/DVN/7ZZWIH). To save time, we will not infer the network and build phylogenomic profiles again. Instead, we will use the pre-built profiles to infer the phylogeny. Our goal here is to reproduce the angiosperm phylogeny in Fig. 1D.

```
#----Get data------------------------------------------------------------------
profiles_file <- file.path(tempdir(), "profiles.tar.gz")
if(!file.exists(profiles_file)) {
    download.file(
        url = "https://dataverse.harvard.edu/api/access/datafile/:persistentId?persistentId=doi:10.7910/DVN/7
        destfile = profiles_file
    )
}
system2("tar", args = c("-zxvf", profiles_file))

#----Load file-----------------------------------------------------------------
prof <- read.table("123genome_profiled_allsize", header = TRUE, sep = " ",
                   row.names = 1)
prof <- as.matrix(prof)

prof[1:10, 1:10] # inspect file
##    pmu ppe pbr Mald Rchi fve roc Dryd Tori Pand
## 1   55  62  83  144   97  52   9   31  101  123
## 2    6   6   8   12    6   6   6    6    6    5
## 3    6   6  10   13    6   7   4    6   12   10
## 4   14  13  19   13    9   8   7    9    9   10
## 5    4   4   4    7   12   8   3    3    3    3
## 6    3   3   5    6    3   3   3    2    3    3
## 7    9  10  20   15   17  17   6    5    7    7
## 8    3   3   4    7    3   2   3    4    2    3
## 9    7   7  16   14    6   6   4    6    7    4
## 10  10  10  18    8    8   4   3    5    5    3

#----Binarize and transpose profiles-------------------------------------------
transposed_profiles <- binarize_and_transpose(prof)
transposed_profiles[1:10, 1:10]
##      1 2 3 4 5 6 7 8 9 10
## pmu  1 1 1 1 1 1 1 1 1  1
## ppe  1 1 1 1 1 1 1 1 1  1
## pbr  1 1 1 1 1 1 1 1 1  1
## Mald 1 1 1 1 1 1 1 1 1  1
## Rchi 1 1 1 1 1 1 1 1 1  1
## fve  1 1 1 1 1 1 1 1 1  1
## roc  1 1 1 1 1 1 1 1 1  1
## Dryd 1 1 1 1 1 1 1 1 1  1
## Tori 1 1 1 1 1 1 1 1 1  1
## Pand 1 1 1 1 1 1 1 1 1  1

#----Infer microsynteny-based phylogeny----------------------------------------
## Using Amborella trichopoda as an outgroup
phylo <- infer_microsynteny_phylogeny(transposed_profiles, outgroup = "atr")

## Read tree file
angiosperm_phylogeny <- treeio::read.tree(phylo[10])

## Replace abbreviations with species names
```

```
angiosperm_phylogeny$tip.label
##   [1] "pmu"  "ppe"  "pbr"  "Mald" "Rchi" "fve"  "roc"  "Dryd" "Tori" "Pand"
##  [11] "Mnot" "Zjuj" "csa"  "cme"  "cla"  "Cuma" "Datg" "Begf" "vra"  "van"
##  [21] "pvu"  "gma"  "cca"  "tpr"  "mtr"  "car"  "lja"  "Anan" "Lang" "adu"
##  [31] "Bpen" "Cgla" "Cill" "Qrob" "cru"  "Csat" "Alyr" "ath"  "Bost" "Lmey"
##  [41] "bol"  "bnp"  "bra"  "spa"  "thh"  "tsa"  "Alp"  "aar"  "cgy"  "tha"
##  [51] "cpa"  "Goba" "Ghir" "gra"  "Dzib" "tca"  "Cmax" "csi"  "Xsor" "mes"
##  [61] "rco"  "ptr"  "lus"  "egr"  "Pgra" "spe"  "sly"  "stu"  "Caba" "Cach"
##  [71] "can"  "pax"  "Inil" "Cuca" "coc"  "sin"  "mgu"  "Oeur" "Lsat" "HanX"
##  [81] "dca"  "ach"  "Aeri" "Cqui" "bvu"  "Ahyp" "Mole" "Kalf" "vvi"  "nnu"
##  [91] "Psom" "Mcor" "Aqco" "sbi"  "Sacc" "Zmay" "sit"  "Sevi" "Ecru" "oth"
## [101] "ogl"  "osa"  "oru"  "Opun" "lpe"  "Trdc" "HORV" "bdi"  "aco"  "mac"
## [111] "egu"  "Pdac" "peq"  "Ashe" "Aoff" "Xvis" "spo"  "zom"  "Peam" "CKAN"
## [121] "Lchi" "atr"  "Nymp"
angiosperm_phylogeny$tip.label <- stringr::str_replace_all(
    angiosperm_phylogeny$tip.label,
    c("^pmu" = "Prunus mume",
      "^ppe" = "Prunus persica",
      "^pbr" = "Pyrus x bretschneideri",
      "^Mald" = "Malus domestica",
      "^Rchi" = "Rosa chinensis",
      "^fve" = "Fragaria vesca",
      "^roc" = "Rubus occidentalis",
      "^Dryd" = "Dryas drummondii",
      "^Pand" = "Parasponia andersonii",
      "^Tori" = "Trema orientale",
      "^Mnot" = "Morus notabilis",
      "^Zjuj" = "Ziziphus jujuba",
      "^cme" = "Cucumis melo",
      "^csa" = "Cucumis sativus",
      "^cla" = "Citrullus lanatus",
      "^Cuma" = "Cucurbita maxima",
      "^Begf" = "Begonia fuchsioides",
      "^Datg" = "Datisca glomerata",
      "^van" = "Vigna angularis",
      "^vra" = "Vigna radiata",
      "^pvu" = "Phaseolus vulgaris",
      "^gma" = "Glycine max",
      "^cca" = "Cajanus cajan",
      "^mtr" = "Medicago truncatula",
      "^tpr" = "Trifolium pratense",
      "^car" = "Cicer arietinum",
      "^lja" = "Lotus japonicus",
      "^Lang" = "Lupinus angustifolius",
      "^Anan" = "Ammopiptanthus nanus",
      "^adu" = "Arachis duranensis",
      "^Cgla" = "Casuarina glauca",
      "^Bpen" = "Betula pendula",
      "^Cill" = "Carya illinoinensis",
      "^Qrob" = "Quercus robur",
      "^bnp" = "Brassica napus",
```

```
               "^bol" = "Brassica oleracea",
               "^bra" = "Brassica rapa",
               "^spa" = "Schrenkiella parvula",
               "^tsa" = "Thellungiella salsuginea",
               "^thh" = "Thellungiella halophila",
               "^ath" = "Arabidopsis thaliana",
               "^Alyr" = "Arabidopsis lyrata",
               "^Csat" = "Camelina sativa",
               "^cru" = "Capsella rubella",
               "^Bost" = "Boechera stricta",
               "^Lmey" = "Lepidium meyenii",
               "^Alp" = "Arabis alpina",
               "^aar" = "Aethionema arabicum",
               "^tha" = "Tarenaya hassleriana",
               "^cgy" = "Cleome gynandra",
               "^cpa" = "Carica papaya",
               "^Ghir" = "Gossypium hirsutum",
               "^Goba" = "Gossypium barbadense",
               "^gra" = "Gossypium raimondii",
               "^Dzib" = "Durio zibethinus",
               "^tca" = "Theobroma cacao",
               "^csi" = "Citrus sinensis",
               "^Cmax" = "Citrus maxima",
               "^Xsor" = "Xanthoceras sorbifolium",
               "^rco" = "Ricinus communis",
               "^mes" = "Manihot esculenta",
               "^ptr" = "Populus trichocarpa",
               "^lus" = "Linum usitatissimum",
               "^Pgra" = "Punica granatum",
               "^egr" = "Eucalyptus grandis",
               "^Cach" = "Capsicum chinense",
               "^Caba" = "Capsicum baccatum",
               "^can" = "Capsicum annuum",
               "^sly" = "Solanum lycopersicum",
               "^spe" = "Solanum pennellii",
               "^stu" = "Solanum tuberosum",
               "^pax" = "Petunia axillaris",
               "^Cuca" = "Cuscuta campestris",
               "^Inil" = "Ipomoea nil",
               "^coc" = "Coffea canephora",
               "^mgu" = "Mimulus guttatus",
               "^sin" = "Sesamum indicum",
               "^Oeur" = "Olea europaea",
               "^HanX" = "Helianthus annuus",
               "^Lsat" = "Lactuca sativa",
               "^dca" = "Daucus carota",
               "^Aeri" = "Actinidia eriantha",
               "^ach" = "Actinidia chinensis",
               "^bvu" = "Beta vulgaris",
               "^Cqui" = "Chenopodium quinoa",
               "^Ahyp" = "Amaranthus hypochondriacus",
```

```
            "^Kalf" = "Kalanchoe fedtschenkoi",
            "^Mole" = "Malania oleifera",
            "^vvi" = "Vitis vinifera",
            "^nnu" = "Nelumbo nucifera",
            "^Mcor" = "Macleaya cordata",
            "^Psom" = "Papaver somniferum",
            "^Aqco" = "Aquilegia coerulea",
            "^Sevi" = "Setaria viridis",
            "^sit" = "Setaria italica",
            "^Ecru" = "Echinochloa crus-galli",
            "^Sacc" = "Saccharum officinarum",
            "^sbi" = "Sorghum bicolor",
            "^Zmay" = "Zea mays",
            "^oth" = "Oropetium thomaeum",
            "^osa" = "Oryza sativa",
            "^ogl" = "Oryza glaberrima",
            "^oru" = "Oryza rufipogon",
            "^Opun" = "Oryza punctata",
            "^lpe" = "Leersia perrieri",
            "^HORV" = "Hordeum vulgare",
            "^Trdc" = "Triticum turgidum",
            "^bdi" = "Brachypodium distachyon",
            "^aco" = "Ananas comosus",
            "^mac" = "Musa acuminata",
            "^Pdac" = "Phoenix dactylifera",
            "^egu" = "Elaeis guineensis",
            "^Ashe" = "Apostasia shenzhenica",
            "^peq" = "Phalaenopsis equestris",
            "^Aoff" = "Asparagus officinalis",
            "^Xvis" = "Xerophyta viscosa",
            "^zom" = "Zostera marina",
            "^spo" = "Spirodela polyrhiza",
            "^CKAN" = "Cinnamomum kanehirae",
            "^Peam" = "Persea americana",
            "^Lchi" = "Liriodendron chinense",
            "^Nymp" = "Nymphaea colorata",
            "^atr" = "Amborella trichopoda"
            )
)

#----Plot tree-----------------------------------------------------------------
suppressPackageStartupMessages(library(ggtree))
ggtree(angiosperm_phylogeny) +
    geom_tiplab(size = 3) +
    xlim(0, 0.3)
```

**Supplementary Material - syntenet: an R/Bioconductor package for the inference and analysis of synteny networks**

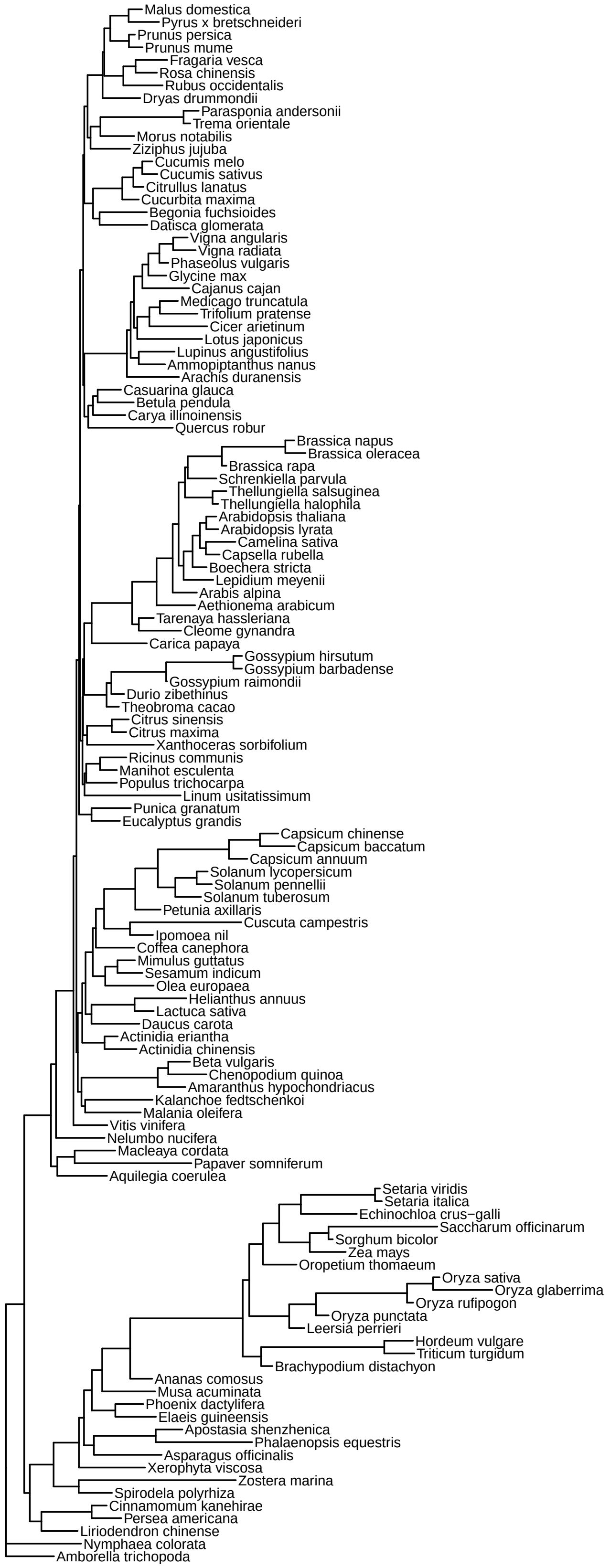

### Section 3: Exploring synteny networks for Chlorophyta genomes

In this final section, we will demonstrate the whole pipeline using the genomes of algae from the Chlorophyta clade.

First, let load the data on-the-fly from Pico-Plaza 3.0 (Van Bel et al. 2018).

```
#----Get proteomes-------------------------------------------------------------
proteome_urls <- c(
    Aprotothecoides = "ftp://ftp.psb.ugent.be/pub/plaza/plaza_pico_03/Fasta/proteome.selected_transcript.apr
    Helicosporidiumsp = "ftp://ftp.psb.ugent.be/pub/plaza/plaza_pico_03/Fasta/proteome.selected_transcript.hs
    Chlorellasp = "ftp://ftp.psb.ugent.be/pub/plaza/plaza_pico_03/Fasta/proteome.selected_transcript.cnc64a.
    PicRCC4223 = "ftp://ftp.psb.ugent.be/pub/plaza/plaza_pico_03/Fasta/proteome.selected_transcript.prcc4223
    PicSE3 = "ftp://ftp.psb.ugent.be/pub/plaza/plaza_pico_03/Fasta/proteome.selected_transcript.pse3.fasta.g
    Asterochlorissp = "ftp://ftp.psb.ugent.be/pub/plaza/plaza_pico_03/Fasta/proteome.selected_transcript.acg
    Csubellipsoidea = "ftp://ftp.psb.ugent.be/pub/plaza/plaza_pico_03/Fasta/proteome.selected_transcript.cvu
    Creinhardtii = "ftp://ftp.psb.ugent.be/pub/plaza/plaza_pico_03/Fasta/proteome.selected_transcript.cre.fas
    Vcarteri = "ftp://ftp.psb.ugent.be/pub/plaza/plaza_pico_03/Fasta/proteome.selected_transcript.vca.fasta.
    Bprasinos = "ftp://ftp.psb.ugent.be/pub/plaza/plaza_pico_03/Fasta/proteome.selected_transcript.bprrcc110
    Otauri = "ftp://ftp.psb.ugent.be/pub/plaza/plaza_pico_03/Fasta/proteome.selected_transcript.ota.fasta.gz
    Osp = "ftp://ftp.psb.ugent.be/pub/plaza/plaza_pico_03/Fasta/proteome.selected_transcript.orcc809.fasta.g
    Olucimarinus = "ftp://ftp.psb.ugent.be/pub/plaza/plaza_pico_03/Fasta/proteome.selected_transcript.olu.fas
    Omediterraneus = "ftp://ftp.psb.ugent.be/pub/plaza/plaza_pico_03/Fasta/proteome.selected_transcript.ome.
    Msp = "ftp://ftp.psb.ugent.be/pub/plaza/plaza_pico_03/Fasta/proteome.selected_transcript.mrcc299.fasta.g
    Mpusilla = "ftp://ftp.psb.ugent.be/pub/plaza/plaza_pico_03/Fasta/proteome.selected_transcript.mpu.fasta.
    Ppatens = "ftp://ftp.psb.ugent.be/pub/plaza/plaza_pico_03/Fasta/proteome.selected_transcript.ppa.fasta.g
)

## Headers in .fa files have protein IDs. Read proteomes and keep only gene IDs
proteomes <- lapply(proteome_urls, function(x) {
    seq <- Biostrings::readAAStringSet(x)
    names(seq) <- gsub(".* \\| ", "", names(seq))
    return(seq)
})

#----Get gene ranges-----------------------------------------------------------
granges_urls <- c(
    Aprotothecoides = "ftp://ftp.psb.ugent.be/pub/plaza/plaza_pico_03/GFF/apr/annotation.selected_transcript
    Helicosporidiumsp = "ftp://ftp.psb.ugent.be/pub/plaza/plaza_pico_03/GFF/hsp/annotation.selected_transcri
    Chlorellasp = "ftp://ftp.psb.ugent.be/pub/plaza/plaza_pico_03/GFF/cnc64a/annotation.selected_transcript.
    PicRCC4223 = "ftp://ftp.psb.ugent.be/pub/plaza/plaza_pico_03/GFF/prcc4223/annotation.selected_transcript
    PicSE3 = "ftp://ftp.psb.ugent.be/pub/plaza/plaza_pico_03/GFF/pse3/annotation.selected_transcript.exon_fe
    Asterochlorissp = "ftp://ftp.psb.ugent.be/pub/plaza/plaza_pico_03/GFF/acg/annotation.selected_transcript
    Csubellipsoidea = "ftp://ftp.psb.ugent.be/pub/plaza/plaza_pico_03/GFF/cvu/annotation.selected_transcript
    Creinhardtii = "ftp://ftp.psb.ugent.be/pub/plaza/plaza_pico_03/GFF/cre/annotation.selected_transcript.ex
    Vcarteri = "ftp://ftp.psb.ugent.be/pub/plaza/plaza_pico_03/GFF/vca/annotation.selected_transcript.exon_f
    Bprasinos = "ftp://ftp.psb.ugent.be/pub/plaza/plaza_pico_03/GFF/bprrcc1105/annotation.selected_transcrip
    Otauri = "ftp://ftp.psb.ugent.be/pub/plaza/plaza_pico_03/GFF/ota/annotation.selected_transcript.exon_fea
    Osp = "ftp://ftp.psb.ugent.be/pub/plaza/plaza_pico_03/GFF/orcc809/annotation.selected_transcript.exon_fe
    Olucimarinus = "ftp://ftp.psb.ugent.be/pub/plaza/plaza_pico_03/GFF/olu/annotation.selected_transcript.ex
```

```
    Omediterraneus = "ftp://ftp.psb.ugent.be/pub/plaza/plaza_pico_03/GFF/ome/annotation.selected_transcript.e
    Msp = "ftp://ftp.psb.ugent.be/pub/plaza/plaza_pico_03/GFF/mrcc299/annotation.selected_transcript.exon_fea
    Mpusilla = "ftp://ftp.psb.ugent.be/pub/plaza/plaza_pico_03/GFF/mpu/annotation.selected_transcript.exon_fe
    Ppatens = "ftp://ftp.psb.ugent.be/pub/plaza/plaza_pico_03/GFF/ppa/annotation.selected_transcript.exon_fea
)

## Read files and keep only required features and columns
annotation <- lapply(granges_urls, function(x) {
    dfile <- file.path(tempdir(), basename(x))
    utils::download.file(x, dfile)
    ranges <- rtracklayer::import(dfile)
    unlink(dfile)

    ranges <- ranges[ranges$type == "gene", ]
    ranges$Parent <- NULL
    ranges$phase <- NULL
    ranges$score <- NULL
    ranges$source <- NULL
    ranges$pid <- NULL
    return(ranges)
})
```

Now, let's proceed to the analyses.

```
# Inspect data
head(proteomes, 3)
## $Aprotothecoides
## AAStringSet object of length 6810:
##        width seq                                          names
##    [1]   177 MEGVHRQLLETTTQWNGDQVAP...LYGLIVGIILASKAGTATPI* AP00G00010
##    [2]   514 MLAVLARRTGGARFPLSSLNLV...GGVGKEMAMMRRELRRRLLG* AP00G00020
##    [3]   416 MEAETAQPGDETPGTTAHMRDP...VDGWCTAERRSKRRSWKAKS* AP00G00030
##    [4]   362 MEALSGTRASFAGATHAFTARK...KRQAKITTELSEIVAGASSV* AP00G00040
##    [5]   310 MSFVTVGGEASSRPTFFELVAA...CPVTLEPVKLEDIWRLYPGM* AP00G00050
##    ...   ... ...
## [6806]  1099 MAGFAADPQVELPPIQYAWLKK...EPLIRGVLARGKATQAAATR* AP00G68060
## [6807]   208 MGRGSRGPRPRLFAKGKCQADG...TGPPPPACLLMVPLAAVSVQ* AP00G68070
## [6808]   254 MATTPQAKTLTRLIQPSLSWLR...HDAEPLGNSYSDAEYVPRRR* AP00G68080
## [6809]   339 MGDPRDVACKLHIMESPAQDWV...DDLRISRSRGGIRVFLRGQA* AP00G68090
## [6810]   220 MARRKRFRSPVLRGPRPRALRH...EALASCASPSRRRRTARWKG* AP00G68100
##
## $Helicosporidiumsp
## AAStringSet object of length 6033:
##        width seq                                          names
##    [1]   172 RSASSKEAGRSATSKEAGRSAT...VRKRPEATCRVASGPAWDTRG HE0000G00010
##    [2]   353 MSTEGHPRYKEVRKIGKGAYGV...EQMLRTASEADAHMPPDHKK* HE0000G00020
##    [3]   239 MVARWKRSKTAGLRSGLELVHA...QPWVVTGGADGTLALFVDDP* HE0000G00030
##    [4]    70 MALWRRKKCSILPPAWMSIANL...LFTGDRDEALAASVFGQHVTR HE0000G00040
##    [5]   311 VLDEIQMIGDQARGWAWTRALL...LAPASTRDPRIAAALLRFAAR HE0000G00050
##    ...   ... ...
## [6029]   112 MLLGLRLWGTIQTSYGGVLQGL...FGAMRALESFAQLLWGEDADG HE0000G60290
```

```
## [6030]   157 LYLGGFNSQEDAARAYDLAALA...EQSSIDPCVELRPLLRLNNDP HE0000G60300
## [6031]   549 SLYTNPRLGRGAGRGAAERVAA...GEAHSQFAKSADWRRRGASRL HE0000G60310
## [6032]   400 MPSLKLRIKPTSGSPPFDVEAS...ANMRAMMQMTEAMQQLQGGPM HE0000G60320
## [6033]   194 MKIWAEVGAPGSKRVPATSKAK...APVKNVPRGAGGKGSKSTGR* HE0000G60330
##
## $Chlorellasp
## AAStringSet object of length 9485:
##        width seq                                          names
##    [1]   449 MQALQLSSRATFAVGRPSSARA...AAGLSAEQMAFLERKRSGGR* CNC64A_001G00010
##    [2]   286 MAAAAVDATRSTTSSAGGGAGA...REHRRKFQWGVSHAMVWRRK* CNC64A_001G00020
##    [3]   348 MEGSAPRVRSQSIGSMSQLLLA...QCAAALQALADCRRSMAGGP* CNC64A_001G00030
##    [4]  1104 MLRKLSSALSLERPVSPRIHPA...AGHDFRAPVSWAGFRELTEK* CNC64A_001G00040
##    [5]   536 MVGQIIDSSGAHPAEGIPKGMD...RELHDFLTLPAQDIIIAMGL* CNC64A_001G00050
##    ...   ... ...
## [9481]   576 MVATGPMPAAPGMAARTAAAPM...AVGDPSLRQEVELMNRILAA* CNC64A_043G00060
## [9482]  1600 MGTGKRSSYEFNEEEDAHEEID...KPAAKSKAGGARPQLKRARA* CNC64A_043G00070
## [9483]  1046 MAGDDDDFVAGGSEEEEEEYQE...QPPAPAGEAGDGAPPPPPPA* CNC64A_043G00080
## [9484]   345 QDVENFRENVKQGYLPLPTDVT...PNPNDTALNAAGTITQVGPA* CNC64A_043G00090
## [9485]   493 MEEHRQRSGTNELPKLLGSWDL...YVRFGAWMVAGLAVYALYGVH CNC64A_043G00100
head(annotation, 3)
## $Aprotothecoides
## GRanges object with 6810 ranges and 2 metadata columns:
##                        seqnames      ranges strand |     type     gene_id
##                           <Rle>   <IRanges>  <Rle> | <factor> <character>
##      [1] contig_KL662078-KL66.. 10089-11286      + |     gene  AP00G46350
##      [2] contig_KL662078-KL66.. 11827-12024      - |     gene  AP00G46360
##      [3] contig_KL662078-KL66.. 12792-14704      + |     gene  AP00G46370
##      [4] contig_KL662078-KL66.. 14870-15959      + |     gene  AP00G46380
##      [5] contig_KL662078-KL66.. 16290-17247      - |     gene  AP00G46390
##      ...                    ...         ...    ... .      ...         ...
##   [6806] contig_KL662078-KL66..   3893-5588      + |     gene  AP00G37130
##   [6807] contig_KL662078-KL66..   6927-9713      - |     gene  AP00G37140
##   [6808] contig_KL662078-KL66.. 10572-13225      + |     gene  AP00G37150
##   [6809] contig_KL662078-KL66.. 16961-20407      + |     gene  AP00G37160
##   [6810] contig_KL662078-KL66.. 20818-39494      - |     gene  AP00G37170
##   -------
##   seqinfo: 102 sequences from an unspecified genome; no seqlengths
##
## $Helicosporidiumsp
## GRanges object with 6033 ranges and 2 metadata columns:
##                    seqnames      ranges strand |     type      gene_id
##                       <Rle>   <IRanges>  <Rle> | <factor>  <character>
##      [1]   contig_pln.dat.1   1046-2301      - |     gene HE0000G55830
##      [2]   contig_pln.dat.1   3782-4823      - |     gene HE0000G55840
##      [3]   contig_pln.dat.1   7169-9142      - |     gene HE0000G55850
##      [4]   contig_pln.dat.1  9601-11866      - |     gene HE0000G55860
##      [5]   contig_pln.dat.1 12235-12901      - |     gene HE0000G55870
##      ...                ...         ...    ... .      ...          ...
##   [6029] contig_pln.dat.996       1-361      + |     gene HE0000G19640
##   [6030] contig_pln.dat.996    933-1902      - |     gene HE0000G19650
##   [6031] contig_pln.dat.997    931-2406      - |     gene HE0000G03620
```

```
##   [6032] contig_pln.dat.998    300-1973      - |     gene HE0000G21590
##   [6033] contig_pln.dat.999      1-1991      - |     gene HE0000G02310
##   -------
##   seqinfo: 4114 sequences from an unspecified genome; no seqlengths
##
## $Chlorellasp
## GRanges object with 9485 ranges and 2 metadata columns:
##            seqnames          ranges strand |     type          gene_id
##               <Rle>       <IRanges>  <Rle> | <factor>      <character>
##      [1] scaffold_1       3183-7075      - |     gene CNC64A_001G00010
##      [2] scaffold_1       7517-9741      - |     gene CNC64A_001G00020
##      [3] scaffold_1     10118-11528      + |     gene CNC64A_001G00030
##      [4] scaffold_1     12061-20718      + |     gene CNC64A_001G00040
##      [5] scaffold_1     21478-27209      - |     gene CNC64A_001G00050
##      ...        ...             ...    ... .      ...              ...
##   [9481] scaffold_9 1593853-1597308      + |     gene CNC64A_009G03300
##   [9482] scaffold_9 1597360-1606723      - |     gene CNC64A_009G03310
##   [9483] scaffold_9 1607086-1609119      + |     gene CNC64A_009G03320
##   [9484] scaffold_9 1609850-1611856      - |     gene CNC64A_009G03330
##   [9485] scaffold_9 1612172-1623917      + |     gene CNC64A_009G03340
##   -------
##   seqinfo: 42 sequences from an unspecified genome; no seqlengths

#----Data processing------------------------------------------------------------
## Check if input objects satisfy required conditions to enter the pipeline
check_input(proteomes, annotation)
## [1] TRUE

## Process the data
pdata <- process_input(proteomes, annotation)

#----Infer synteny network------------------------------------------------------
## Run DIAMOND
diamond <- run_diamond(seq = pdata$seq)

## Network inference per se
start_time <- Sys.time() # not required, just to get starting time
algae_network <- infer_syntenet(
    blast_list = diamond,
    annotation = pdata$annotation
)
end_time <- Sys.time() # not required, just to get end time

head(algae_network) # inspect data
##                  Anchor1               Anchor2
## 1 Chlo_CNC64A_028G00030 Chlo_CNC64A_035G00060
## 2 Chlo_CNC64A_028G00040 Chlo_CNC64A_035G00070
## 3 Chlo_CNC64A_028G00070 Chlo_CNC64A_035G00080
## 4 Chlo_CNC64A_028G00080 Chlo_CNC64A_035G00090
## 5 Chlo_CNC64A_028G00110 Chlo_CNC64A_035G00140
## 6 Chlo_CNC64A_028G00140 Chlo_CNC64A_035G00150
```

Now, we have the synteny network for Chlorophyta algae. Let's see how long it took to infer the network.

```
end_time - start_time
## Time difference of 53.29748 secs
```

Finally, let's explore phylogenomic profiles for this network and infer a microsynteny-based phylogeny for these algae genomes. We will use *Physcomitrium patens* as outgroup for the tree.

```
#----Network clustering and phylogenomic profiling------------------------------
## Get synteny clusters
clusters <- cluster_network(algae_network)
head(clusters) # inspect data
##                      Gene Cluster
## 1 Chlo_CNC64A_028G00030       1
## 2 Chlo_CNC64A_028G00040       2
## 3 Chlo_CNC64A_028G00070       3
## 4 Chlo_CNC64A_028G00080       4
## 5 Chlo_CNC64A_028G00110       5
## 6 Chlo_CNC64A_028G00140       6
```

```
## Get phylogenomic profiles
profiles <- phylogenomic_profile(clusters)
```

```
head(profiles$profile_matrix) # inspect data
##
##          Apro Aste Bpra Chlo Crei Csub Mpus Msp Oluc Omed Osp Otau PicR PicS
##   12323    0    0    0    0    1    0    0   0    0    0   0    0    0    0
##   12322    0    0    0    0    1    0    0   0    0    0   0    0    0    0
##   12321    0    0    0    0    1    0    0   0    0    0   0    0    0    0
##   12320    0    0    0    0    1    0    0   0    0    0   0    0    0    0
##   12319    0    0    0    0    1    0    0   0    0    0   0    0    0    0
##   12318    0    0    0    0    1    0    0   0    0    0   0    0    0    0
##
##          Ppat Vcar
##   12323    0    1
##   12322    0    1
##   12321    0    1
##   12320    0    1
##   12319    0    1
##   12318    0    1

#----Plot profiles--------------------------------------------------------------
## Create a data frame with species names and abbreviations in colnames() of
## profiles matrix
species_order <- data.frame(
    Species = c(
        "Aprotothecoides", "Chlorellasp", "PicRCC4223",
        "PicSE3", "Asterochlorissp", "Csubellipsoidea", "Creinhardtii",
        "Vcarteri", "Bprasinos", "Otauri", "Osp", "Olucimarinus",
        "Omediterraneus", "Msp", "Mpusilla", "Ppatens"
```

```
        ),
        Abbrev = c(
            "Apro", "Chlo", "PicR", "PicS", "Aste", "Csub", "Crei",
            "Vcar", "Bpra", "Otau", "Osp", "Oluc", "Omed", "Msp",
            "Mpus", "Ppat"
        )
    )

    ## Add taxonomic information
    species_order$Clade <- c(
        rep("Trebouxiophyceae", 6),
        rep("Chlamydomonadales", 2),
        rep("Marmiellales", 7),
        "Outgroup"
    )

    ## Reorder columns and add longer names
    profiles$profile_matrix <- profiles$profile_matrix[, species_order$Abbrev]
    colnames(profiles$profile_matrix) <- species_order$Species

    ## Plotting heatmap
    plot_profiles(
        profiles,
        species_annotation = species_order[, c("Species", "Clade")],
        cluster_species = species_order$Species
    )
```

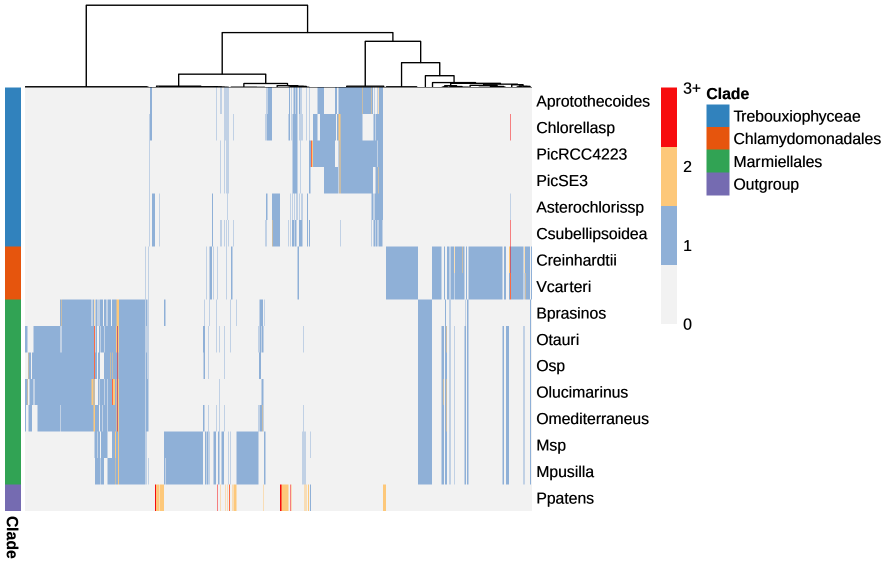

We can see that the phylogenomic profiles of synteny clusters match the species' phylogeny. For instance, there are Chlamydomonadales-specific clusters, Marmiellales-specific clusters, and Trebouxiophyceae-specific clusters. Besides, the phylogenomic profiles of algae is very different from that of *Physcomitrium patens*.

Now, let's use these profiles to infer a microsynteny-based phylogeny. Here, as the phylogenomic profiles do not contain constant sites, we will add `+ASC` to the model to perform an ascertainment bias correction.

```
#----Infer microsynteny-based phylogeny----------------------------------------
## Binarize and tranpose profiles matrix
bt_mat <- binarize_and_transpose(profiles$profile_matrix)
bt_mat[1:5, 1:5] # inspect data
##
##                   12323 12322 12321 12320 12319
##   Aprotothecoides     0     0     0     0     0
##   Chlorellasp         0     0     0     0     0
##   PicRCC4223          0     0     0     0     0
##   PicSE3              0     0     0     0     0
##   Asterochlorissp     0     0     0     0     0

## Infer phylogeny using P. patens as outgroup
phylo <- infer_microsynteny_phylogeny(
    bt_mat,
    outgroup = "Ppatens",
    model = "MK+ASC+R",
    threads = 1
)

## Plot tree
suppressPackageStartupMessages(library(ggtree))
algae_tree <- treeio::read.tree(phylo[10])
ggtree(algae_tree) +
    geom_tiplab(size = 3) +
    xlim(0, 0.2)
```

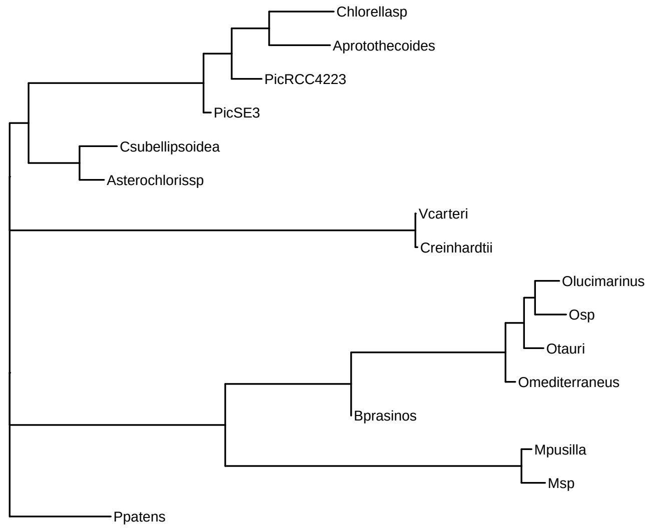

Overall, the microsynteny-based tree is in line with the alignment-based tree (see https://bioinformatics.psb.ugent.be/plaza/versions/plaza_pico_03/). However, the tree topology shows Picochlorum_RCC4223 closer to the Chlorella sp-Auxenochlorella protothecoides clade than to Picochlorum sp. SENEW3 (SE3). The phylogenetic relationships could be better resolved if more genomes are included.

### Session information

This document was created under the following conditions.

```
## - Session info ---------------------------------------------------------------
##  setting  value
##  version  R version 4.2.1 (2022-06-23)
##  os       Ubuntu 20.04.4 LTS
##  system   x86_64, linux-gnu
##  ui       X11
##  language (EN)
##  collate  en_US.UTF-8
##  ctype    en_US.UTF-8
##  tz       Europe/Brussels
##  date     2022-08-19
##  pandoc   2.18 @ /usr/lib/rstudio/bin/quarto/bin/tools/ (via rmarkdown)
##
## - Packages -------------------------------------------------------------------
##  package          * version  date (UTC) lib source
##  ape                5.6-2    2022-03-02 [1] CRAN (R 4.2.0)
##  aplot              0.1.6    2022-06-03 [1] CRAN (R 4.2.0)
##  assertthat         0.2.1    2019-03-21 [1] CRAN (R 4.2.0)
##  backports          1.4.1    2021-12-13 [1] CRAN (R 4.2.0)
##  BiocGenerics       0.42.0   2022-04-26 [1] Bioconductor
##  BiocManager        1.30.18  2022-05-18 [1] CRAN (R 4.2.0)
##  BiocStyle        * 2.25.0   2022-06-15 [1] Github (Bioconductor/BiocStyle@7150c28)
##  Biostrings         2.64.0   2022-04-26 [1] Bioconductor
##  bitops             1.0-7    2021-04-24 [1] CRAN (R 4.2.0)
##  bookdown           0.27     2022-06-14 [1] CRAN (R 4.2.0)
##  broom              0.8.0    2022-04-13 [1] CRAN (R 4.2.0)
##  cellranger         1.1.0    2016-07-27 [1] CRAN (R 4.2.0)
##  cli                3.3.0    2022-04-25 [1] CRAN (R 4.2.0)
##  cluster            2.1.3    2022-03-28 [1] CRAN (R 4.2.0)
##  coda               0.19-4   2020-09-30 [1] CRAN (R 4.2.0)
##  codetools          0.2-18   2020-11-04 [1] CRAN (R 4.2.0)
##  colorspace         2.0-3    2022-02-21 [1] CRAN (R 4.2.0)
##  crayon             1.5.1    2022-03-26 [1] CRAN (R 4.2.0)
##  DBI                1.1.3    2022-06-18 [1] CRAN (R 4.2.0)
##  dbplyr             2.2.1    2022-06-27 [1] CRAN (R 4.2.1)
##  digest             0.6.29   2021-12-01 [1] CRAN (R 4.2.0)
##  dplyr            * 1.0.9    2022-04-28 [1] CRAN (R 4.2.0)
##  ellipsis           0.3.2    2021-04-29 [1] CRAN (R 4.2.0)
##  evaluate           0.15     2022-02-18 [1] CRAN (R 4.2.0)
##  fansi              1.0.3    2022-03-24 [1] CRAN (R 4.2.0)
```

```
##  farver             2.1.0    2021-02-28 [1] CRAN (R 4.2.0)
##  fastmap            1.1.0    2021-01-25 [1] CRAN (R 4.2.0)
##  forcats          * 0.5.1    2021-01-27 [1] CRAN (R 4.2.0)
##  fs                 1.5.2    2021-12-08 [1] CRAN (R 4.2.0)
##  generics           0.1.2    2022-01-31 [1] CRAN (R 4.2.0)
##  GenomeInfoDb       1.32.2   2022-05-15 [1] Bioconductor
##  GenomeInfoDbData   1.2.8    2022-05-06 [1] Bioconductor
##  GenomicRanges      1.48.0   2022-04-26 [1] Bioconductor
##  ggfun              0.0.6    2022-04-01 [1] CRAN (R 4.2.0)
##  ggnetwork          0.5.10   2021-07-06 [1] CRAN (R 4.2.0)
##  ggplot2          * 3.3.6    2022-05-03 [1] CRAN (R 4.2.0)
##  ggplotify          0.1.0    2021-09-02 [1] CRAN (R 4.2.0)
##  ggtree           * 3.4.0    2022-04-26 [1] Bioconductor
##  glue               1.6.2    2022-02-24 [1] CRAN (R 4.2.0)
##  gridGraphics       0.5-1    2020-12-13 [1] CRAN (R 4.2.0)
##  gtable             0.3.0    2019-03-25 [1] CRAN (R 4.2.0)
##  haven              2.5.0    2022-04-15 [1] CRAN (R 4.2.0)
##  hms                1.1.1    2021-09-26 [1] CRAN (R 4.2.0)
##  htmltools          0.5.2    2021-08-25 [1] CRAN (R 4.2.0)
##  htmlwidgets        1.5.4    2021-09-08 [1] CRAN (R 4.2.0)
##  httr               1.4.3    2022-05-04 [1] CRAN (R 4.2.0)
##  igraph             1.3.2    2022-06-13 [1] CRAN (R 4.2.0)
##  intergraph         2.0-2    2016-12-05 [1] CRAN (R 4.2.0)
##  IRanges            2.30.0   2022-04-26 [1] Bioconductor
##  jsonlite           1.8.0    2022-02-22 [1] CRAN (R 4.2.0)
##  knitr              1.39     2022-04-26 [1] CRAN (R 4.2.0)
##  labeling           0.4.2    2020-10-20 [1] CRAN (R 4.2.0)
##  lattice            0.20-45  2021-09-22 [1] CRAN (R 4.2.0)
##  lazyeval           0.2.2    2019-03-15 [1] CRAN (R 4.2.0)
##  lifecycle          1.0.1    2021-09-24 [1] CRAN (R 4.2.0)
##  lubridate          1.8.0    2021-10-07 [1] CRAN (R 4.2.0)
##  magrittr           2.0.3    2022-03-30 [1] CRAN (R 4.2.0)
##  MASS               7.3-57   2022-04-22 [1] CRAN (R 4.2.0)
##  Matrix             1.4-1    2022-03-23 [1] CRAN (R 4.2.0)
##  mgcv               1.8-40   2022-03-29 [1] CRAN (R 4.2.0)
##  modelr             0.1.8    2020-05-19 [1] CRAN (R 4.2.0)
##  munsell            0.5.0    2018-06-12 [1] CRAN (R 4.2.0)
##  network            1.17.2   2022-05-21 [1] CRAN (R 4.2.0)
##  networkD3          0.4      2017-03-18 [1] CRAN (R 4.2.0)
##  nlme               3.1-158  2022-06-15 [1] CRAN (R 4.2.0)
##  patchwork          1.1.1    2020-12-17 [1] CRAN (R 4.2.0)
##  permute            0.9-7    2022-01-27 [1] CRAN (R 4.2.0)
##  pheatmap           1.0.12   2019-01-04 [1] CRAN (R 4.2.0)
##  pillar             1.7.0    2022-02-01 [1] CRAN (R 4.2.0)
##  pkgconfig          2.0.3    2019-09-22 [1] CRAN (R 4.2.0)
##  purrr            * 0.3.4    2020-04-17 [1] CRAN (R 4.2.0)
##  R6                 2.5.1    2021-08-19 [1] CRAN (R 4.2.0)
##  RColorBrewer       1.1-3    2022-04-03 [1] CRAN (R 4.2.0)
##  Rcpp               1.0.8.3  2022-03-17 [1] CRAN (R 4.2.0)
##  RCurl              1.98-1.7 2022-06-09 [1] CRAN (R 4.2.0)
##  readr            * 2.1.2    2022-01-30 [1] CRAN (R 4.2.0)
```

```
##  readxl            1.4.0    2022-03-28 [1] CRAN (R 4.2.0)
##  reprex            2.0.1    2021-08-05 [1] CRAN (R 4.2.0)
##  rlang             1.0.3    2022-06-27 [1] CRAN (R 4.2.1)
##  rmarkdown         2.14     2022-04-25 [1] CRAN (R 4.2.0)
##  rstudioapi        0.13     2020-11-12 [1] CRAN (R 4.2.0)
##  rvest             1.0.2    2021-10-16 [1] CRAN (R 4.2.0)
##  S4Vectors         0.34.0   2022-04-26 [1] Bioconductor
##  scales            1.2.0    2022-04-13 [1] CRAN (R 4.2.0)
##  sessioninfo       1.2.2    2021-12-06 [1] CRAN (R 4.2.0)
##  statnet.common    4.6.0    2022-05-02 [1] CRAN (R 4.2.0)
##  stringi           1.7.6    2021-11-29 [1] CRAN (R 4.2.0)
##  stringr         * 1.4.0    2019-02-10 [1] CRAN (R 4.2.0)
##  syntenet        * 0.99.4   2022-08-13 [1] Github (almeidasilvaf/syntenet@2963c7f)
##  tibble          * 3.1.7    2022-05-03 [1] CRAN (R 4.2.0)
##  tidyr           * 1.2.0    2022-02-01 [1] CRAN (R 4.2.0)
##  tidyselect        1.1.2    2022-02-21 [1] CRAN (R 4.2.0)
##  tidytree          0.3.9    2022-03-04 [1] CRAN (R 4.2.0)
##  tidyverse       * 1.3.1    2021-04-15 [1] CRAN (R 4.2.0)
##  treeio            1.20.0   2022-04-26 [1] Bioconductor
##  tzdb              0.3.0    2022-03-28 [1] CRAN (R 4.2.0)
##  utf8              1.2.2    2021-07-24 [1] CRAN (R 4.2.0)
##  vctrs             0.4.1    2022-04-13 [1] CRAN (R 4.2.0)
##  vegan             2.6-2    2022-04-17 [1] CRAN (R 4.2.0)
##  withr             2.5.0    2022-03-03 [1] CRAN (R 4.2.0)
##  xfun              0.31     2022-05-10 [1] CRAN (R 4.2.0)
##  xml2              1.3.3    2021-11-30 [1] CRAN (R 4.2.0)
##  XVector           0.36.0   2022-04-26 [1] Bioconductor
##  yaml              2.3.5    2022-02-21 [1] CRAN (R 4.2.0)
##  yulab.utils       0.0.4    2021-10-09 [1] CRAN (R 4.2.0)
##  zlibbioc          1.42.0   2022-04-26 [1] Bioconductor
##
##  [1] /home/faalm/R/x86_64-pc-linux-gnu-library/4.2
##  [2] /usr/local/lib/R/site-library
##  [3] /usr/lib/R/site-library
##  [4] /usr/lib/R/library
##
## ------------------------------------------------------------------------------
```
